## Supplementary Figures for "TMC7 deficiency causes acrosome biogenesis defects and male infertility in mice"

### Supplementary Information

#### Figure S1

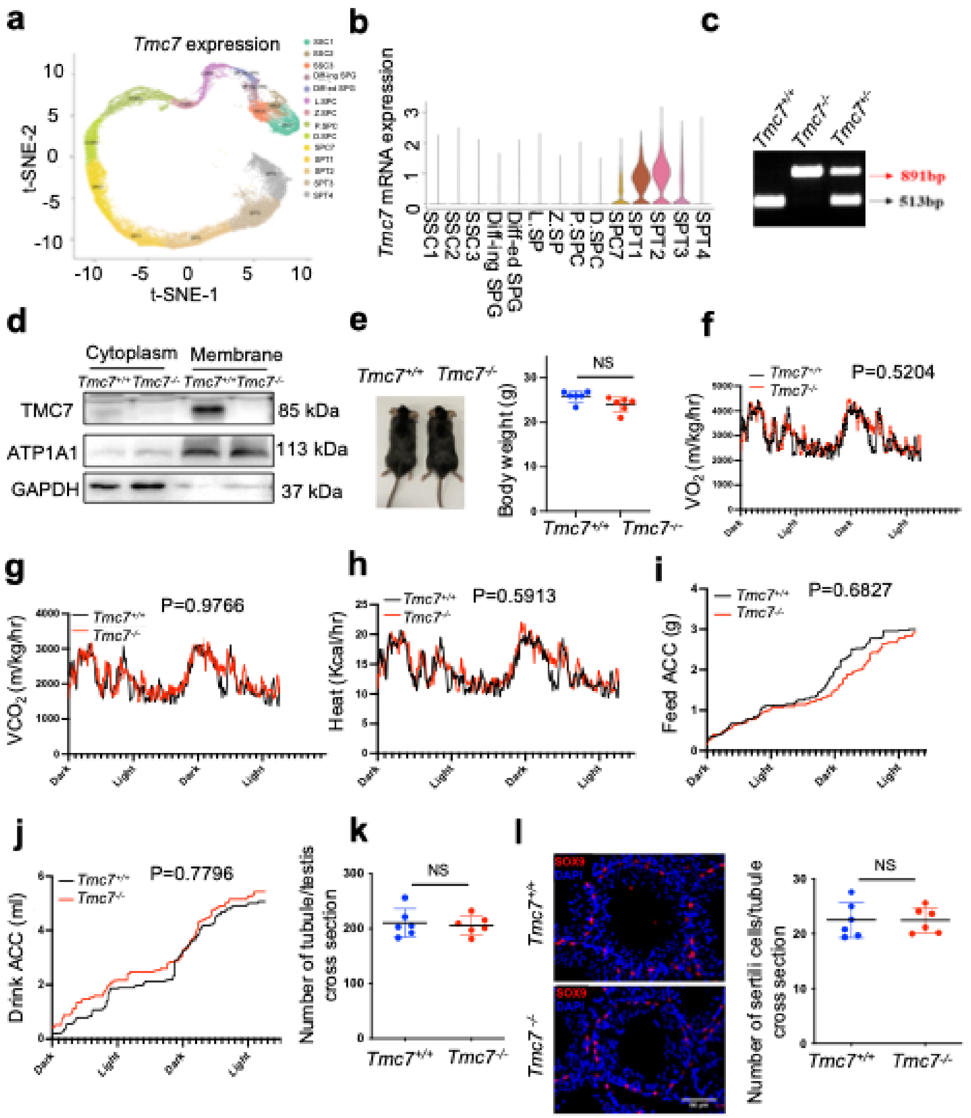


**Fig S1. Phenotypic characterization of *Tmc7*$^{-/-}$ mice.**

(a, b) T-Distributed Stochastic Neighbor Embedding(t-SNE) plot and violin plot showed the expression of TMC7 in different human germ cell using Male Health Atlas（http://malehealthatlas.cn/). SSC1-3: Spermatogonial stem cells I-III; Diff-ed SPG: Differenced spermatogonia; Diff-ing SPG: differentiating spermatogonia; SPC: spermatocyte; SPT: spermatids. (c) genotyping of *Tmc7* mice by PCR of tail-derived DNA. (d) Western blot was performed to detect the knockout efficiency in cytoplasm and membrane protein of testis in 9-Week mice. (e) The mice of WT and *Tmc7*$^{-/-}$ for 9-Week mice and the body weights of 9-Week WT and *Tmc7*$^{-/-}$ male mice, Six mice of each genotype were used. (f-j) Oxygen and consumption, carbon dioxide production, energy expenditure, food and water intake were measured using metabolic cages in 9-week WT and *Tmc7*$^{-/-}$ male mice. Four mice were used. two-way analysis of variance (ANOVA) was used to analyze the P-values. (k) The average number of tubules per testis in 9-week WT and *Tmc7*$^{-/-}$ male mice, six mice were used. (l) IF staining of SOX9, a marker of sertoli cell and average number of Sertoli cells per seminiferous tubules in 9-week WT and *Tmc7*$^{-/-}$ male mice.10 tubules in each testis were counted and six mice were used. Scale bar, 50μm. The results are presented as the mean ± S.D. Two-sided unpaired Student's t-test and two-way analysis of variance (ANOVA) were used to analyze the P-values. (ns, stands for non-significant, *P < 0.05, **P < 0.01, ***P < 0.001).

**Figure S2**

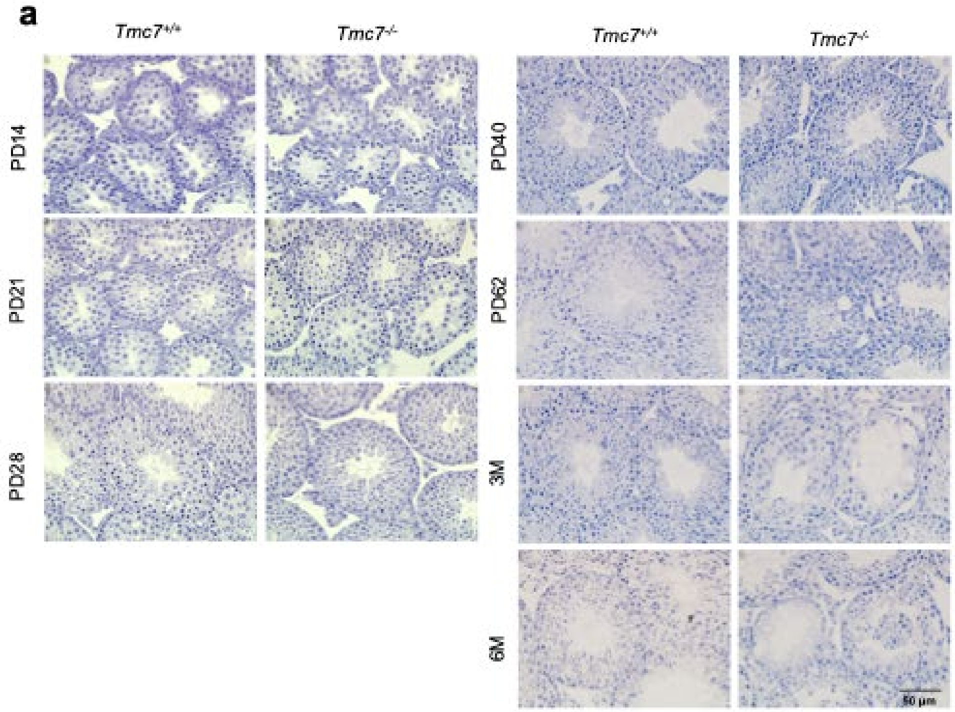


**Fig S2. Histological detection in WT and *Tmc7*$^{-/-}$ mice.**

(a) Hematoxylin staining in testis of WT and *Tmc7*$^{-/-}$ mice at different postnatal day points (PD14, PD21, PD28, PD40, PD62, 3M and 6M). Scale bar, 50μm.

**Figure S3**

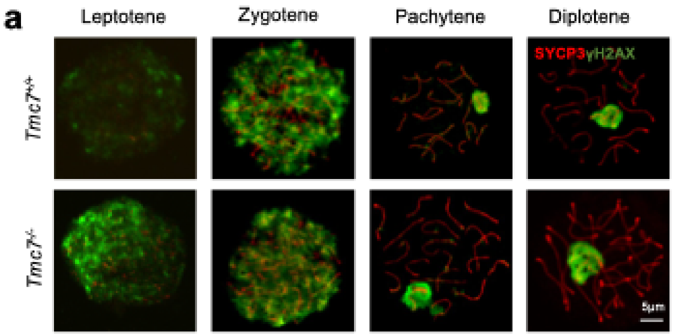


**Fig S3. TMC7 is not required for meiotic progression.**

(a) IF staining with antibody against SYCP3(red) and γ-H2AX (green), the marker of meiotic in chromosome spreads of spermatocytes from the testes in PD20 WT and *Tmc7*$^{-/-}$ mice. Scale bar, 5μm.

**Figure S4**

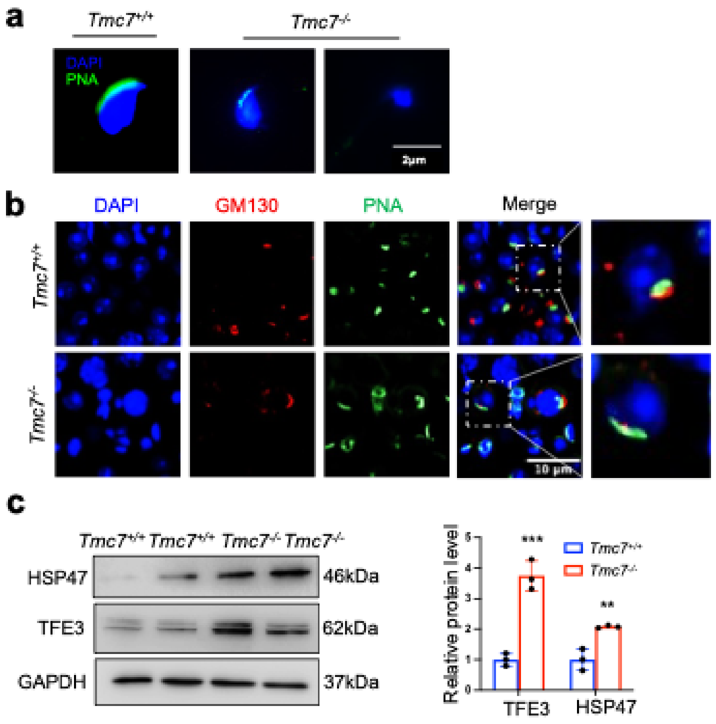


**Fig S4. The defects of acrosome and Golgi apparatus in *Tmc7*$^{-/-}$ spermatozoa.**

(a) Acrosome staining by using fluorescein-conjugated peanut agglutinin (PNA), a protein that binds to the outer acrosomal membrane in sperm collected from caudal epididymis. Scale bar, 2μm. (b) IF staining of GM130 (red) and PNA (green) in squashing tubules of 9-Week WT and *Tmc7*$^{-/-}$ mice. Nuclei were stained with DAPI (blue). Scale bar, 10μm. (c) Western blot analysis of protein levels of Golgi stress-associated proteins (HSP47, TFE3) in 9-week-old WT and *Tmc7*$^{-/-}$ testes. Image J was used to quantify the protein level. Images are representative of at least three independent experiments. The results are presented as the mean ± S.D., and two-sided unpaired Student's t-tests were used to calculate the P-values. (*P < 0.05, **P < 0.01, ***P < 0.001).

**Figure S5**

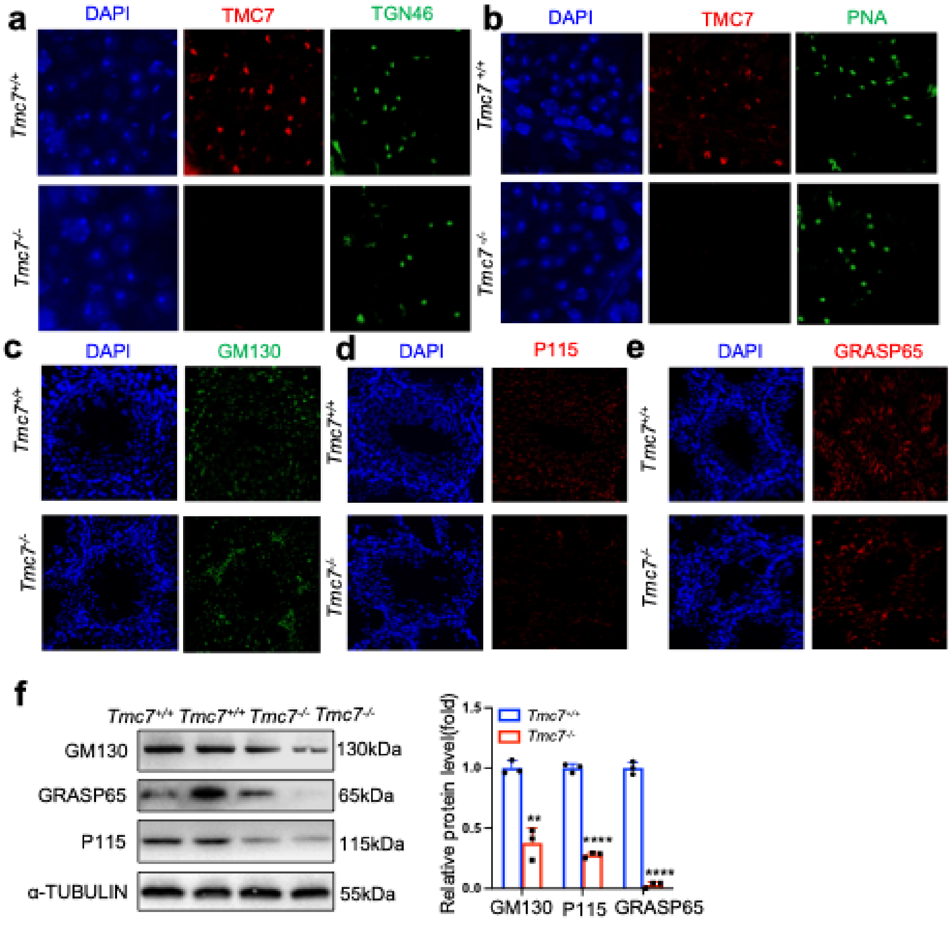


**Fig S5. The location of TMC7 in testis.**

(a) IF staining of TMC7 (red) and TGN46 (green), DAPI indicates the nucleus in seminiferous tubule. (b) IF staining of TMC7 (red) and PNA (green), DAPI indicates the nucleus in seminiferous tubule. (c) IF staining GM130 (green), DAPI indicates the nucleus in seminiferous tubule. (d) IF staining P115 (red), DAPI indicates the nucleus in seminiferous tubule. (e) IF staining GRASP65 ((red), DAPI indicates the nucleus in seminiferous tubule. (f) Western blots of the protein levels of GM130 and the GM130-interacting proteins P115 and GRASP65 in 9-week-old WT and *Tmc7*$^{-/-}$ testes. Image J was used to quantify the protein level. Images are representative of at least three independent experiments. The results are presented as the mean

± S.D., and two-sided unpaired Student's t-tests were used to calculate the P-values. (*P < 0.05, **P < 0.01, ***P < 0.001).

**Figure S6**

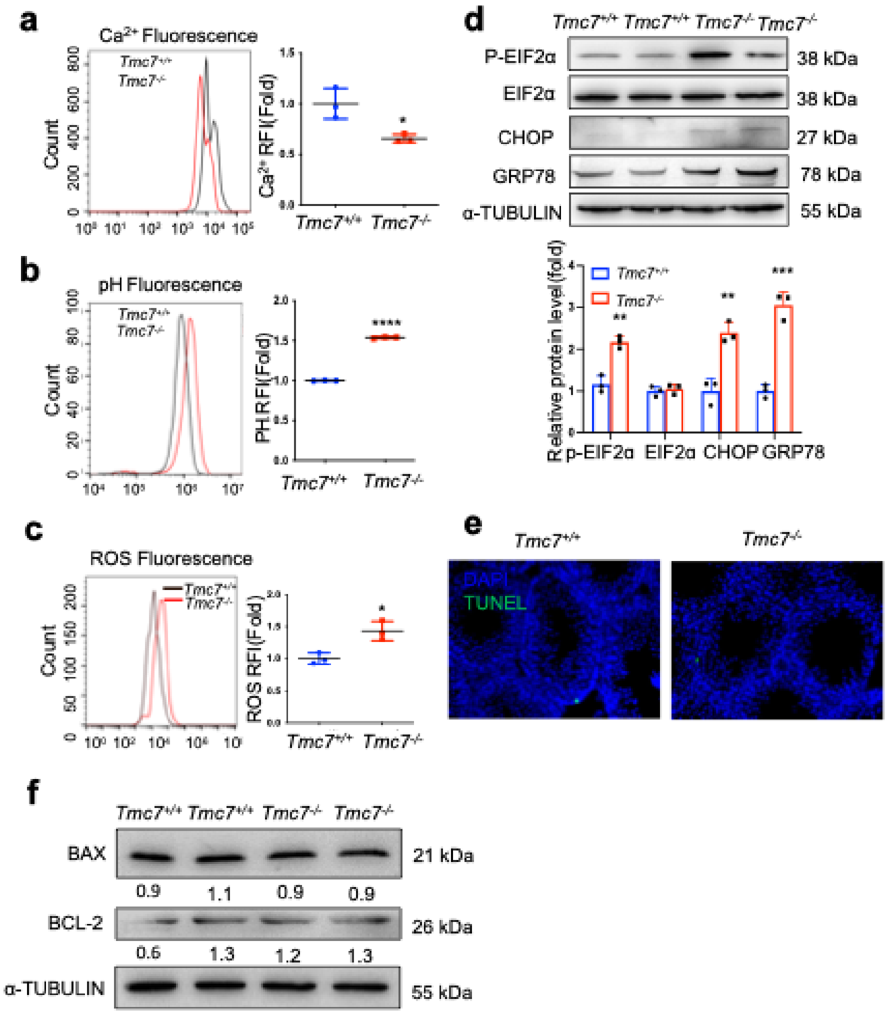


**Fig S6. Intracellular homeostasis was impaired in *Tmc7*$^{-/-}$ testis.**

(a) The $Ca^{2+}$ levels were measured by flow cytometry in WT and *Tmc7*$^{-/-}$ germ cells isolated from PD30 testes. There mice were used. (b) The pH levels were measured by flow cytometry in WT and *Tmc7*$^{-/-}$ germ cells isolated from PD30 testes. There mice were used. (c) The ROS levels were measured by flow cytometry in WT and *Tmc7*$^{-/-}$ germ cells isolated from PD30 testes. There mice were used. (d) Western blot analysis of protein levels of ER chaperone-associated proteins (p-EIF2α, EIF2α, Chop, and GRP78) in 9-week-old WT and *Tmc7*$^{-/-}$ testes. Image J was used to quantify the protein level. Images are representative of at least three independent experiments. (e) TUNEL assay of PD30 WT and *Tmc7*$^{-/-}$ male mice. (f) Apoptosisrelated proteins (BAX, BCL2) were detected by western blotting in PD30 WT and *Tmc7*$^{-/-}$ testes. Image J was used to quantify the protein level. The results are presented as the mean ± S.D., and two-sided unpaired Student's t-tests were used to calculate the P-values. (*P < 0.05, **P < 0.01, ***P < 0.001).

**Figure S7**

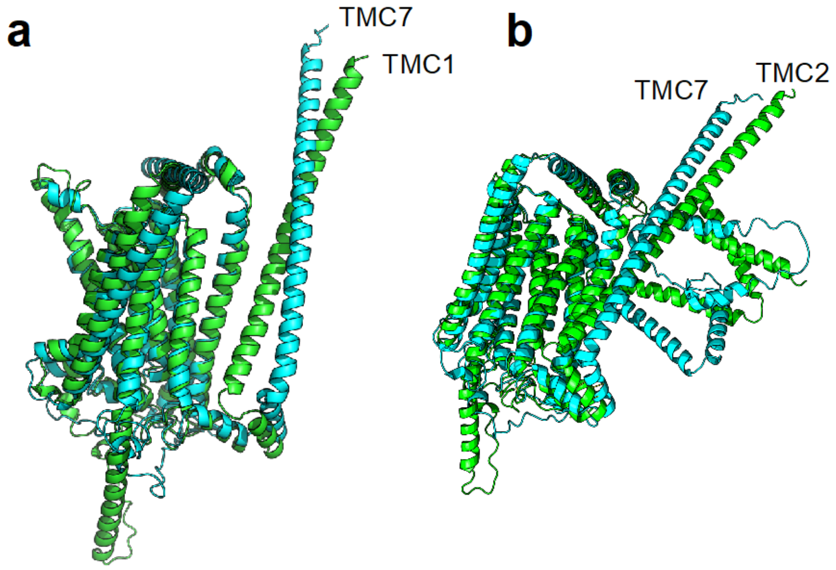


**Fig S7. Structure comparison of TMC1/2 with TMC7.**

(a) The structure of TMC1 with TMC7. (b) The structure of TMC2 with TMC7.

**Figure S8**

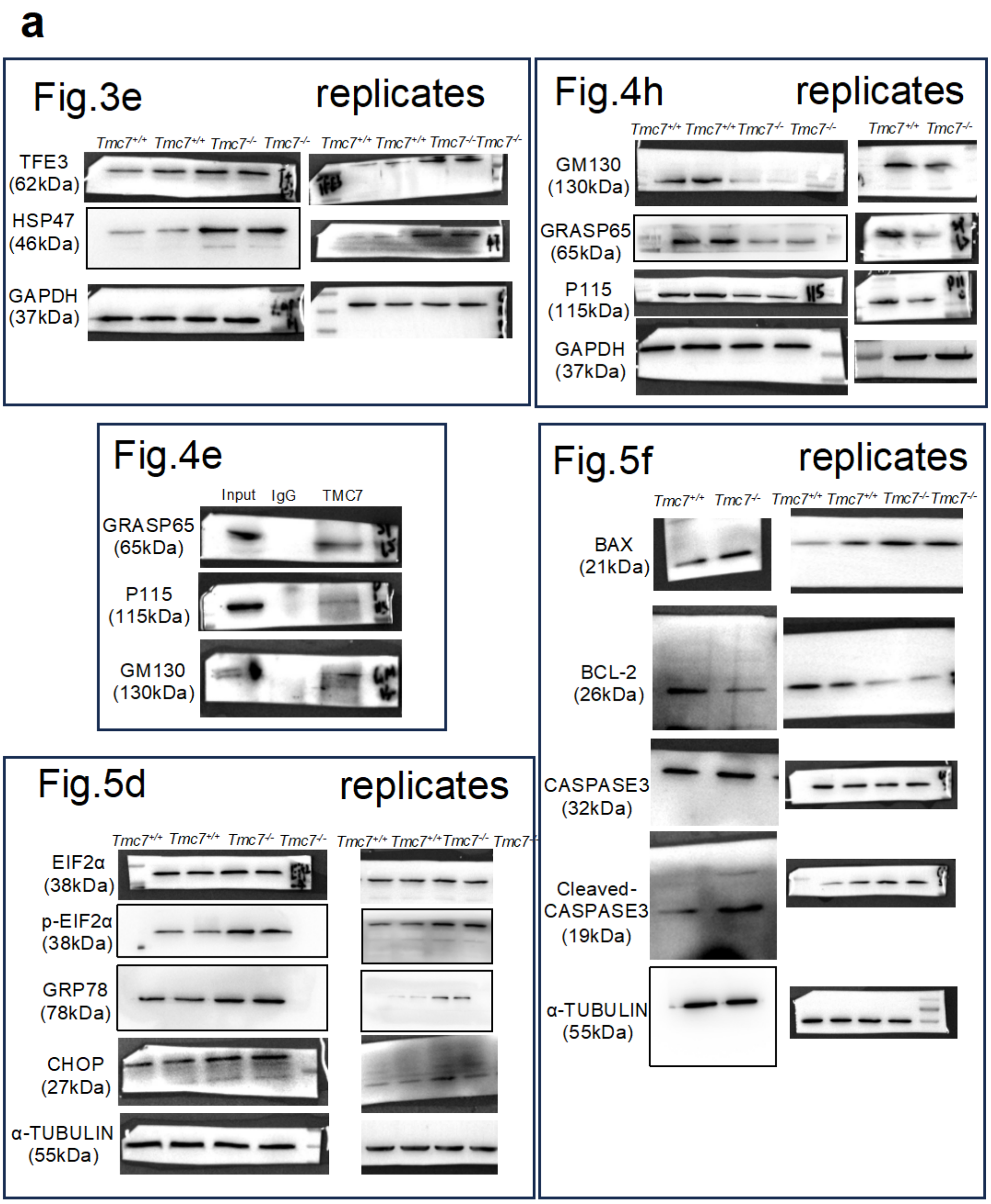
a
Fig.3e
replicates
Tmc7+/+ Tmc7+/+ Tmc7-/- Tmc7-/-
Tmc7+/+ Tmc7+/+ Tmc7-/- Tmc7-/-
TFE3 (62kDa)
HSP47 (46kDa)
GAPDH (37kDa)
Fig.4h
replicates
Tmc7+/+ Tmc7+/+ Tmc7-/- Tmc7-/-
Tmc7+/+ Tmc7-/-
GM130 (130kDa)
GRASP65 (65kDa)
P115 (115kDa)
GAPDH (37kDa)
Fig.4e
Input IgG TMC7
GRASP65 (65kDa)
P115 (115kDa)
GM130 (130kDa)
Fig.5f
replicates
Tmc7+/+ Tmc7-/-
Tmc7+/+ Tmc7+/+ Tmc7-/- Tmc7-/-
BAX (21kDa)
BCL-2 (26kDa)
CASPASE3 (32kDa)
Cleaved-CASPASE3 (19kDa)
α-TUBULIN (55kDa)
Fig.5d
replicates
Tmc7+/+ Tmc7+/+ Tmc7-/- Tmc7-/-
Tmc7+/+ Tmc7+/+ Tmc7-/- Tmc7-/-
EIF2α (38kDa)
p-EIF2α (38kDa)
GRP78 (78kDa)
CHOP (27kDa)
α-TUBULIN (55kDa)

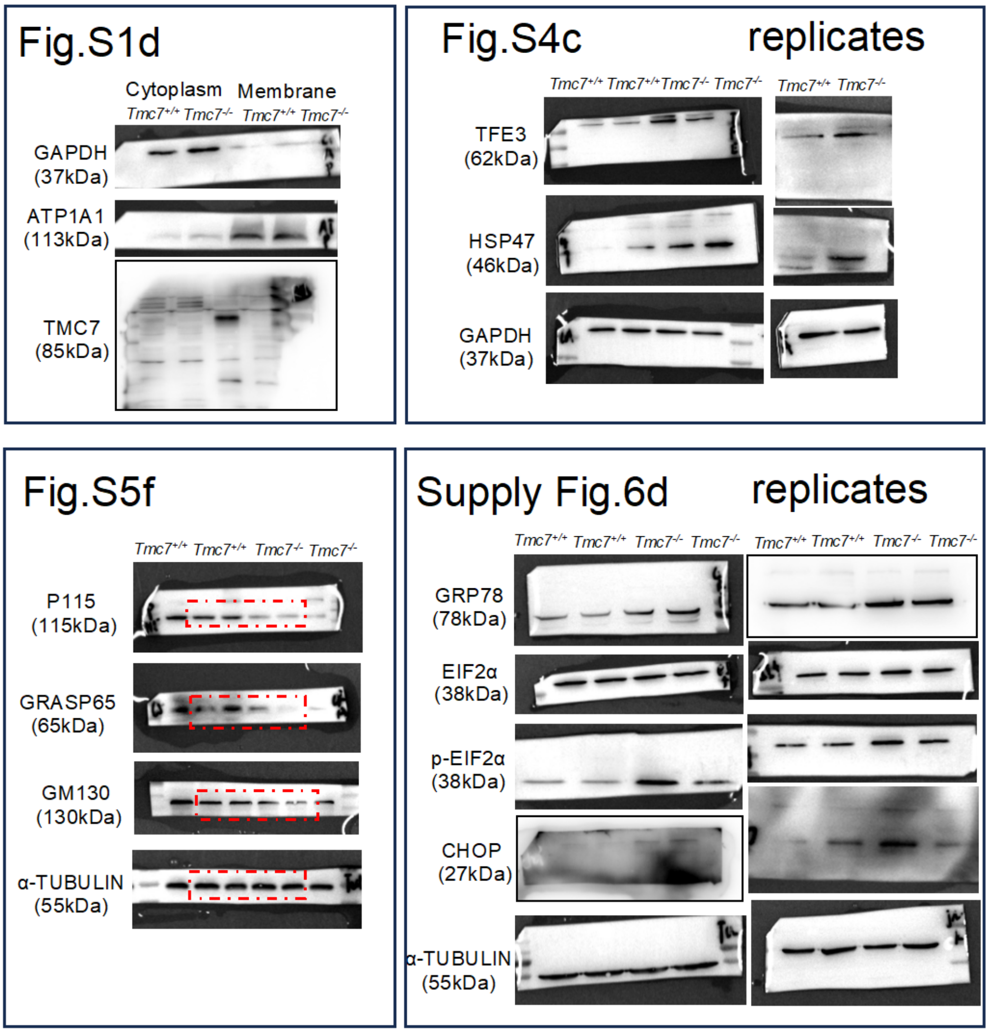


**Fig S8. Raw data of western blot**

(a) Raw data of western blot and replicates.
