## Supplementary Table for "TMC7 deficiency causes acrosome biogenesis defects and male infertility in mice"

Table1. Fertility testing of *Tmc7*^+/-^, *Tmc7*^-/-^ male mice and *Tmc7*^-/-^ female mice.

| **Male** | **Female** | **Num of pups/litter** |
| --- | --- | --- |
| *Tmc7^-/-^* | WT * 2 | None |
| *Tmc7^-/-^* | WT * 2 | None |
| *Tmc7^-/-^* | WT * 2 | None |
| *Tmc7^+/-^* | *Tmc7^+/-^* * 2 | 8,5,6,8 |
| *Tmc7^+/-^* | *Tmc7^-/-^* * 2 | 9,10,11,8 |
| *Tmc7^+/-^* | *Tmc7^-/-^* * 2 | 8,6,5,8 |
